# Coarse-grained Martini 3 model of chondroitin sulfate A

**DOI:** 10.1101/2025.10.20.683363

**Authors:** Paulius Greicius, Frauke Gräter, Fabian Grünewald, Camilo Aponte-Santamaría

## Abstract

Chondroitin sulfate A (CSA) is a negatively charged linear glycosaminoglycan which plays a vital role in many biological processes. Research on CSA has been challenging due to its size, chemical heterogeneity, and multitude of binding partners. To address these issues, we developed a model of CSA for coarse-grained molecular dynamics simulations based on the Martini 3 force-field. We demonstrate that this model is capable of reproducing atomistic properties of the repeating CSA disaccharide unit, including its molecular volume and bonded interactions, and structural polymer properties of CSA chains of different lengths. In particular, for biologically-relevant long chains and despite of using an explicit solvent, the computational cost is significantly reduced, relative to the cost equivalent atomistic simulations would require. The compatibility of the model with the Martini Gō protein model was tested by retrieving the force-response of the CSA–malaria adhesin VAR2CSA complex. Importantly, we explored the influence of electrostatics on CSA aggregation. We show that the default Martini 3 parameters lead to over-aggregation. We provide at least three different strategies to alleviate this issue, making use of a bigger bead for sodium cations, reflecting its hydration shell, partial ionic charges, as a mean-field resource to take into account electronic polarizability, and, optionally, particle-mesh Ewald summation as a more robust treatment of long-range electrostatics. Our model opens the door for predictive modeling of CSA and potentially other chondroitin sulfates. In addition, this model provides insights for the further development of coarse-grained models of highly-charged systems.

## Introduction

Chondroitin sulfate (CS) is a linear glycosaminoglycan composed out of alternating N-acetyl galactosamine (GAL) and glucuronic acid (GLA) units connected via *β*1,3 and *β*1,4 linkages, respectively (Figure 1A). Sulfates can be variably added at various hydroxyls of the disaccharide, creating a highly negatively charged sugar. The polysaccharide is synthesized in the endoplasmic reticulum and Golgi where it is also attached to cell surface or secreted proteins via serine residues^1^. CS is a key extracellular matrix component in various tissues including skin^2^, bone^3^, cartilage^4^, vasculature^5^, cornea^6^ and central nervous system^7^. Chains isolated from *in vivo* sources display high heterogeneity both in their molecular weight (50-150 kDa) and frequency of sulfated residues ^8^. Such structural diversity reflects the fact that CS serves a broad range of biological functions.

**Figure 1.**
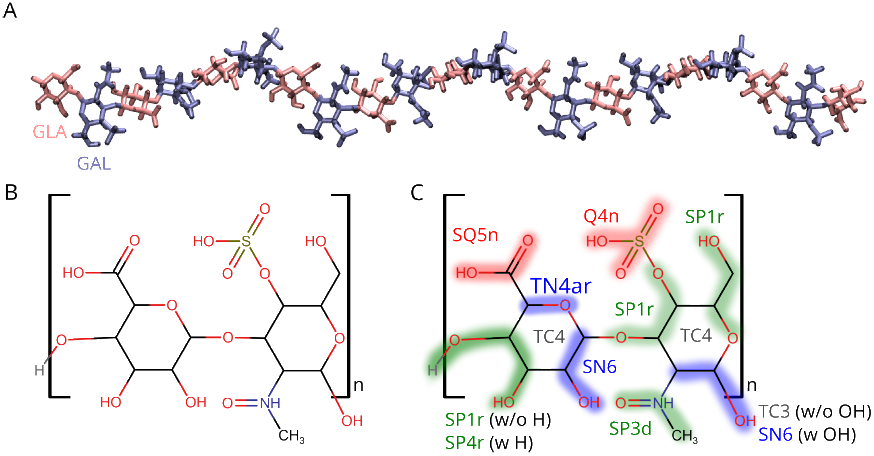
Structure and mapping of CSA into Martini 3 beads. (A) Structure of a CSA 21-mer. D-glucuronic acid (GLA) is colored in pink and the N-acetyl-D-galactosamine 4-sulfate (GAL) in blue. (B) Chemical structure of the GLA-GAL repeating unit. (C) Mapping of CSA repeating unit to Martini 3 beads. Shaded areas indicate atoms belonging to the same bead. Text next to each bead indicates the bead type. TC4 is the virtual site located in the center of the ring. Bead types which are part of the glycosidic bond are labeled with “w/o”, meanwhile beads at the end of the chain are labeled with “w”.

The polyanionic nature and variable sulfation status of chondroitin sulfate chains enable it to form interactions with various cations^9^ or proteins^1,7^. Heavily sulfated CS chains create high osmotic pressure, essential for load bearing in articulate cartilage^10^, sustaining bone toughness^11^ and hydration of cornea^12^. Free carboxylate and sulfate groups chelate metal ions, especially divalent cations such as Calcium^13^. The ability of CS to sequester metal ions enhances its radical scavenging activity^14^ making it an effective antioxidant^15^. Similarly, chelation of extracellular Ca^2+^ affects transmembrane potential, enabling modulation of voltage gated ion channel activity^16^. The sulfation pattern on CS chains also dictates interactions with specific growth factors, morphogens, cytokines or chemokines thereby regulating various signaling pathways^1,17,18^. In the nervous system, CS plays a role in axon growth and guidance, but depending on CS sulfation, it can either promote or inhibit these processes^7,19^. Overall, while chondroitin sulfate has been implicated in many biological phenomena, it remains challenging to delineate specific roles and properties of different CS chain types.

Chondroitin sulfate A (CSA), characterized by sulfate on carbon 4 of GAL residues (Figure 1B) is one of the most common CS types in animal tissues^1^. It is the major type of CS in human brain^20^ or at sites of spinal cord injury^21^. CSA reduces with age in cartilage^4^ and has been associated to bone formation by facilitating collagen mineralization^22^. The sugar is also involved in diseases: as a ligand for placental malaria pathogen^23,24^ and a marker of cancer cells^25^. In addition, CSA has received attention as a dietary supplement against osteoarthritis^26^ and as a component of hydrogels or drug delivery systems^27^. However, research on this sugar is complicated by the heterogeneous nature of the samples^8,28^, sometimes even leading to contradictory results^19^. For this reason, an approach capable of studying well-defined, homogeneous CSA chains is needed to gain accurate insights into the sugar’s properties and interactions.

Molecular dynamics (MD) simulations have tremendously advanced our understanding of glycosaminoglycans, providing key aspects related to their structure, (thermo)dynamics, energetics, and interactions (for a comprehensive reviews see Nagarajan et al^29^ and Perez et al^30^). The technique has the advantage of providing the spatial and temporal evolution of glycan systems at varying level of resolution and with explicitly defined chain chemical composition. In the context of CS, MD has been applied to probe conformations of CS fragments with different sulfation motifs^31,32^ and the conformational impact of cation or hydroxyl radical binding^33–35^. MD and molecular docking have also been employed to examine CS in complex with chemokines CXCL8^36^, CXCL14^37^, malarial protein VAR2CSA^38–40^ and cysteine cathepsins^41,42^. However, all these studies employed CS chains up to 20 monomer units of length due to significant computational demands of all-atom MD simulations. In contrast, chondroitin sulfate length in biological systems commonly exceeds 100 monomers. One way to enable simulations of longer CS chains is using coarse-grained (CG) MD simulations. In coarse-grained approximations, groups of atoms are represented as a single particle (pseudoatom) which reduces the number of entities within the system, thus speeding up its dynamics by reducing total degrees of freedom. CG MD simulations of infinitely diluted chondroitin sulfate, represented by its backbone, has been shown to successfully reproduce experimental measurements including radius of gyration^43^, osmotic pressure^44^, and hydrodynamic radius.^45^ A CG model that also accounts for functional groups was introduced^46^, but significant computational speedup was only achieved in implicit solvent. More recently, a CG model for the most common Glycosaminoglycans, including fully-sulfated CSs, has been developed in the Martini force field (version 2.2.)^47^. Nevertheless, a model for the most common CS form, i.e. CSA, with a single-sulfation at the 4th carbon of GAL, within the framework of generalized coarse-grained force-fields, for large-scale multi-component simulations, is still needed.

Martini 3 is a widely used force-field for coarse-grained simulations of biomolecules with new and improved features over its predecessor Martini 2.2^48^. Models for many carbohydrates were implemented in the force-field^49,50^, but chondroitin sulfate has not yet been defined. In this study we introduce a Martini 3 model for CSA repeating unit which accurately reproduces the behavior of atomistic CHARMM36m^51^ simulations and examine the impact of simulation parameters on CSA aggregation. Then, we proceed to simulate CSA in complex with CSA-binding protein VAR2CSA. Lastly, to illustrate the computational gains of the coarse-grained model, we perform equilibrium simulations of a 123 monomers-long CSA chain.

## Methods

### All-atom Chondroitin sulfate A structure

A linear 21 subunit chondroitin sulfate A all-atom structure and associated GROMACS itp files were generated using the CHARMM-GUI Glycan Modeler^52^. D-glucuronic acid (GLA) subunit was connected to N-acetyl-D-galactosamine (GAL) via beta 1,4 linkage. Then, GAL was connected to GLA via beta 1,3 linkage. The structure was extended in such an alternating fashion until it contained 21 subunits, with 11 GLA and 10 acetyl GAL sugars in total. All N-acetyl-D-galactosamines were modified by attaching O-linked sulfate to the fourth carbon.

### All-atom MD simulations of CSA

For comparison, all-atom MD simulations of either a single 21mer CSA chain or four of them were carried out. The CHARMM36m glycan parameters were used for such simulations^51^. Simulations were carried out in cubic boxes with periodic boundary condition. For the single-chain simulation,the box size was (9.77 nm)^3^ and for the multi-chain simulations (17.36 nm)^3^. In the latter case, the chains were initially positioned such that at least 1 nm of distance was between them. In both single-chain or four-chain simulations, the box was solvated with CHARMM-TIP3P water molecules^53,54^ and neutralized with an excess of sodium ions. In addition, NaCl salt up to a final concentration of 0.15 M was added to the systems. Solvated systems consisted of approximately 90000 atoms (single chain) and 500000 atoms (four chains). Steric clashes were removed by energy minimization via the steepest descent algorithm. Thermalization followed under NVT conditions, during 125 ps of MD at a temperature of 303.15 K. The solvent was subsequently equilibrated around the glycans, by performing 250 ps MD in NPT ensemble, maintaining temperature at 303.15 K and pressure at 1 bar. During minimization, thermalization, and solvent equilibration steps, positional restraints of 400 *kJ/mol/nm*^2^ on backbone heavy atoms of glycan rings, 40 *kJ/mol/nm*^2^ on the side-chain heavy atoms, and 4 *kJ/mol/nm*^2^ dihedral restraints, were imposed. Equilibrium dynamics were then simulated, releasing the positional and dihedral restraints, for 5 replicates of 1 *µ*s each (5 *µ*s cumulative) for each system.

### Coarse-grained simulations of CSA

Coarse-grained simulations of CSA were carried out for a single 21mer, for a system containing four 21mer chains and for a single 123mer. The coarse-grained models for single and four 21mers were generated by center-of-gravity mapping the NPT equilibrated all atom structure, with the Fast-Forward (ff map) tool^55^, retaining the same simulation box dimensions, CSA chain and ion number as well as their positions. 123mer chain was built by extending CG 21mer with PyMol^56^ and solvated in a (51.72 nm)^3^ box with 0.15 M neutralizing salt, resulting in a total of about 1 million particles. The Martini3^48^ coarse-grained force-field was used. NaCl ions were represented as Q5 beads with *±*0.75 charge unless otherwise noted. Martini 3 water (compressing four atomistic water molecules into one bead) was used to solvate the system. Files containing the initial interaction parameters (itp) suitable for GROMACS were generated with Polyply (gen params)^57^. For simulations with non-integer salt ion charges, additional ions were added until the system was fully neutralized. Energy minimization was carried out to remove steric clashes during 2000 steps, with the steepest descent algorithm, followed by equilibration in the NPT ensemble, for 75 ns. Finally, 5 production runs of 1 *µ*s were run for simulations with 21mers and 10 of 2 *µ*s for the 123mer.

### Simulations of VAR2CSA in complex with CSA

Equilibrium coarse-grained MD simulations of the core domain of VAR2CSA (NF54 strain) in complex with a fully sulphated CSA 21mer were carried out. The conformations of the complex were retrieved from previous all-atom equilibrium MD simulations^40^ (at a timepoint 150 ns of each of the 10 replicas). The CSA chain was coarse-grained following the same process as outlined above. The Martini 3 model of VAR2CSA was built using Martinize2 choosing default parameters^58^, including secondary structures predicted by DSSP^59^ and native contacts^60^. In addition, to preserve the tertiary structure of the protein, a Gō model, establishing a network of pair-wise Lennard-Jones interactions of strength *ϵ*_*LJ*_ = 8.414*kJ/mol*, was applied^61,62^. Gō interactions were truncated at a distance lower than 0.3 nm or greater than 1.1 nm and were excluded from the internally disordered region spanning residues from 376 to 551. To remove steric clashes, the complex was energy minimized using steepest descent for 100 steps. The energy-minimized complex was placed in a dodecahedral box, with the periodic boundary walls at a distance of at least 1 nm of the complex and solvated with Martini 3 water. The system was neutralized at an ionic strength of 0.15 M NaCl, both Na and Cl represented as Q5 beads of ±0.75*e* charge, respectively. The non-integer excess in charge was controlled by adding a single TP1 bead with charge *−*0.25*e* which is not expected to introduce any significant interactions in the system. Energy minimization and three 750-ps rounds of NPT equilibration preceded the production runs. During the first round position restraints (with elastic constant of 1000*kJ/mol/nm*^2^) were applied to all protein and glycan beads. During the second round only protein beads were restrained. Finally, the third round was performed without any restraints. Production runs of 300 ns were subsequently performed independently for 10 replicates.

The final conformation after 300 ns of equilibration was used as starting configuration for force-probe MD simulations. Virtual harmonic springs (1000*kJ/mol/nm*^2^ force constant) were attached to the C-terminal methionine 1953 of VAR2CSA and first monomer of CSA 21-mer chain and moved with a constant velocity of 0.1*m/s* in opposite directions. N=10 force-probe MD simulations were carried out, each one 120 ns.

### MD simulation parameters and algorithms

All-atom MD simulations were performed using the GROMACS MD package (version 2025.2) ^63^. Newton’s equations of motion were numerically integrated at time steps of 2 fs (1 fs during thermalization) using the Leap frog algorithm. The temperature (pressure) was kept constant by coupling either simulated system to the Nose-Hoover (Parrinello-Rahman) thermostat (barostat), with coupling constants of 1 ps (5 ps), every 500 (2500) integration steps. Short-range non-bonded van der Waals interactions were modeled by a Lennard Jones potential, truncated at a distance of 1.2 nm. Electrostatic interactions were treated using the particle mesh Ewald (PME) algorithm^64,65^. Neighbors were considered via the Verlet buffer scheme^66^ with a tolerance of 0.005 *kJmol*^*−*1^*ps*^*−*1^. LINCS^67^ was used to constraint the bonds involving heavy atoms of the sugars while SETTLE^68^ was employed to constraint both bonds and angular vibrations of water.

Coarse-grained MARTINI 3 MD simulations were carried out using the same parameters and algorithms as for the all-atom simulations, except for the following differences. The integration timestep was 15 femtoseconds. Temperature pressure coupling was achieved with Berendsen (Berendsen) thermostat (barostat)^69^ during minimization or equilibration and v-rescale^70^ (Parrinello-Rahman^71^) - during production. Temperature coupling constants were 1 ps, 2 ps and 1 ps during minimization, equilibration and production respectively. Meanwhile, the pressure coupling constant was 12 ps during all steps. Instead of a tolerance in the interactions with the Verlet buffer neighbor scheme, a fixed neighbor list cutoff of 1.35 nm was used^72^. Electrostatic interactions were either treated with the reaction-field scheme^73^ or with PME, for the simulations of 21mers and exclusively with PME for the simulations of 123mer or a CSA 21mer in complex with VAR2CSA.

### Analysis

The solvent-accessible surface area (SASA) was determined using the double cubic lattice method as implemented in the GROMACS *gmx sasa* tool^74^ using a probe sphere of size 0.191 nm. Atomic wan der Walls-radii were retrieved from^75^ and for the Martini 3 beads the following sizes were used: 0.264 nm for regular, 0.23 nm for small and 0.191 nm for tiny beads^49^.

Center-of-geometry mapping of the all-atom trajectory to Martini 3 beads and quantification of bonded term distributions was carried out by using Fast-Forward^55^. The bonded term distributions were compared by means of the Jensen-Shannon divergence score, available in the SciPy library^76^. The optimal bin width for each bonded term comparison was selected using the Freedman-Diaconis rule^77^.

GROMACS *gmx polystat* function was used to compute both the end-to-end distance and the radius of gyration of the CSA chains. *gmx distance* was employed for computing pairwise residue distances and to quantify protein-CSA contacts by considering two beads “in contact” if their minimum distance was less than 0.45 nm. The radial distribution function of sodium was calculated using *gmx rdf*, with either the carboxylic acid or sulfate group bead acting as reference particle and sodium bead acting as selection. Root-mean-square fluctuations (RMSF) of the protein residues or CSA monomers was determined with *gmx rmsf* ^63^. Persistence length was defined as the exponential decay half-life of GLA virtual sites autocorrelation function *C*(*x*), quantified with *MDAnalysis*.*analysis*.*polymer*.*PersistenceLength* ^78^. *MDAnalysis*.*analysis*.*rms*.*rmsd* ^79,80^ was used to calculate VAR2CSA root mean square deviation.

The Flory scaling exponent *ν* was determined by fitting the function < *d*_*ij*_ > = *b*|*i − j*|^*ν*^ to the curve < *d*_*ij*_ > versus |*i* − *j*|, where < *d*_*ij*_ > is the average distance between residues *i* and *j* and *b* is the Kuhn length^81^.

For VAR2CSA simulations, analysis was computed for each replicate from 150 ns to 200 ns timepoints in all-atom trajectory and from 100 ns to 300 ns for coarse-grained to account for equilibration. Analysis of CSA 123mer trajectory was done from 500 ns to 3000 ns for the same reason.

Molecular structures were visualized and rendered with VMD^82^.

The CG force-field parameters, MD simulation parameters and initial configurations generated in this study are deposited at the site https://github.com/PaulGreic/CSA_Martini3.

## Results

### Parametrization of the repeating unit

CSA is a linear polysaccharide composed out repeating D-glucuronic acid (GLA) and N-acetyl-D-galactosamine 4-sulfate (GAL) units (Figure 1A,B). We built a Martini3 coarse-grained model of this repeating unit, validating it against all-atom simulations using the CHARMM36m force field. Most of the model was based on existing atom-to-bead definitions from other carbohydrate and small molecule models (Figure 1C)^49,83^. The sulfate group, of key importance for CSA, has not yet been parametrized in Martini3. We mapped it following general force-field guidelines: four heavy atoms, including a period three element sulfur, were represented as a regular size bead of diameter 0.264 nm and Q4n label to account for the -1 charge delocalized among oxygens^48^. In addition, in order to capture the ring stacking interactions as recommended for Martini3 carbohydrates^49^, a virtual interaction site was added to the center of each six-member glycan ring. The resulting model mapped 53 atoms into 11 coarse-grained particles.

As in other Martini3 carbohydrate models, to capture the rigid nature of the glycan rings, beads representing them were connected via distance constraints rather than harmonic bonds^49^. Furthermore, the distance between such beads was increased by 10% compared to atomistic reference, to more accurately represent the molecular volume (Figure S1). This resulted in a distribution of solvent-accessible surface area (SASA) for our Martini 3 model that compared extremely well to that distribution for the atomistic reference (Figure 2A), a result that was visually corroborated by the Connolly surfaces (Figure 2B).Thus, the coarsegrained model adopted here accurately captures the shape of the CSA repeating unit.

**Figure 2.**
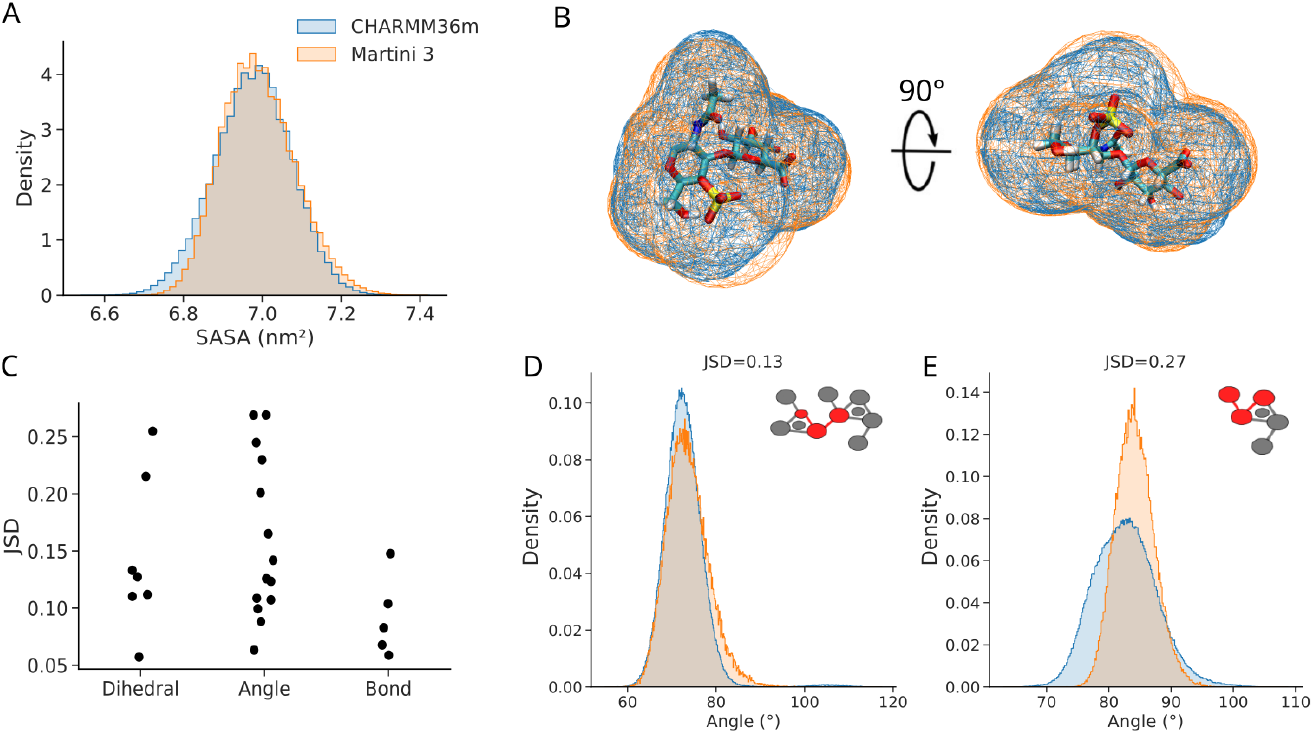
CG Martini 3 model of CSA accurately reproduces its exposure to the solvent and bonded term distributions. (A,B) Solvent-accessible surface area (SASA) distribution (A) and Connolly surface (B) comparison for the GLA-GAL repeating unit of CSA, recovered from MD simulations at all-atom (CHARMM36m) and coarse-grained (Martini 3) resolution. (C) Jensen-Shannon divergence statistic for the three types of bonded terms used, comparing the Martini 3 model with the all-atom CHARMM36m reference data (JSD=0: identical distributions, and JSD large: dissimilar distributions). (D,E) Examples of bonded interaction distributions from coarse-grained and forward-mapped all-atom simulations, for the bonded term marked in red in the inset: one close to the median, representative of a low value of the JSD (D), and one with the largest JSD representing the worst agreement (E). B D and E follow the same color scheme as A. See comparison of distributions for all bonded terms in Figure S2-4.

All other bonded interactions within the disaccharide repeating unit were adjusted to match their corresponding distributions in all-atom simulations. Note that in order to make the comparison possible, all-atom trajectories were forward mapped, based on the center-of-gravity of atom groups corresponding to each Martini bead. The resulting distributions are show in Figures S2–4. The agreement between Martini and reference (all-atom) distributions of each bonded term was quantified using the Jensen-Shannon divergence (JSD), which gives a score of 0 for identical distributions and score of 1 for non-overlapping distributions (Figure 2C).The median JSD value was 0.13, indicating that most of the terms were well reproduced by the coarse-grained model (Figure 2C,D). The worst JSD value (of 0.27) was observed for an angle with asymmetrical reference distribution, a difficult case to reproduce at this level of resolution (Figure 2E). Note that the coarse-grained simulations were run using modified parameters to avoid CSA aggregation (see below), nevertheless similar results were also achieved with default Martini 3 parameters (Figure S5A).

### Evaluation of polymer properties

After establishing a Martini 3 model for the CSA repeating unit, we evaluated whether it is capable of reproducing atomistic polymer properties of a CSA 21mer under equilibrium conditions (Figure 3A). The size of polymers is commonly quantified with the end-to-end distance (*ETE*) or the radius of gyration (*R*_*g*_). For a gaussian chain, ratio 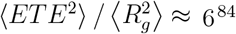. In the reference all-atom simulations the ratio was 10.4 indicating that this glycan chain adopts a highly extended conformation. Reassuringly, a similar level of elongation was observed in our coarse-grained Martini 3 model, where the ratio was estimated to be around10.0 (Figure 3B). Furthermore the characteristic ratio, i.e. 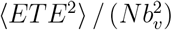 (with *N* = 21, the polymerization degree, and *b*_*v*_ = 0.52 nm, the bond length), was found do be 10.8, which is consistent with the estimate from a previous implicit-solvent CG model of CSA (see value of ∼10 for comparable *N* in Figure 6 in Bathe et al.^43^).

**Figure 3.**
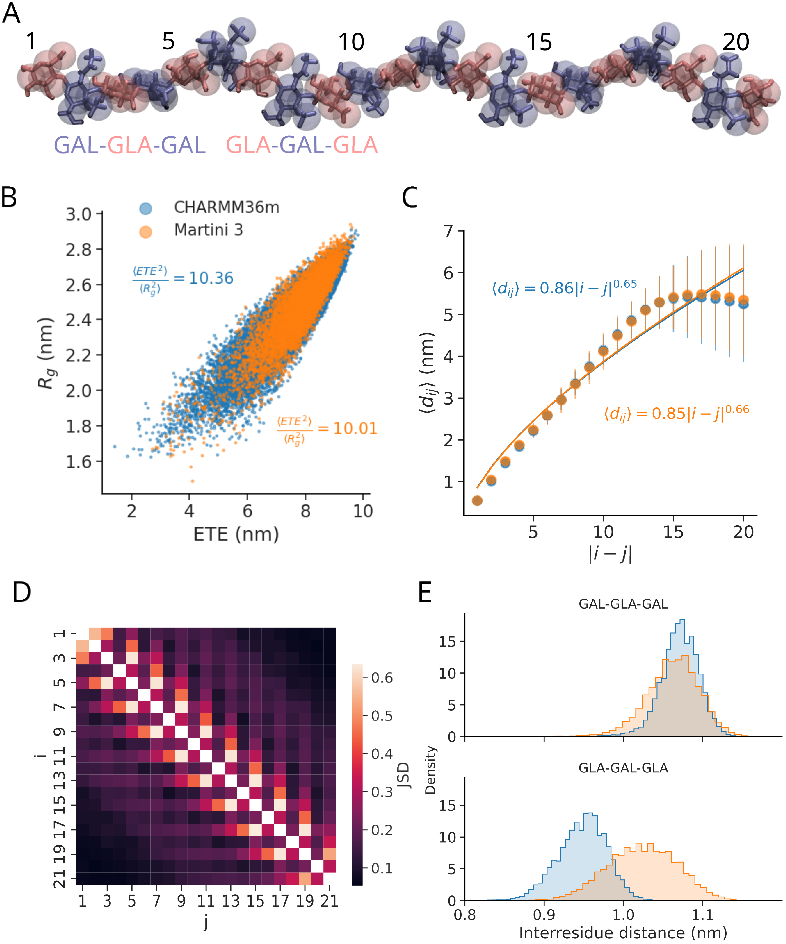
Coarse-grained Martini 3 model captures the polymer behavior of a single CSA chain. (A) A 21-saccharide CSA chain, superposing the all-atom structure (sticks) with the CG Martini 3 model (spheres) highlights the alternating nature of the GAL and GLA units. (B) Ratio between radius of gyration *R*_*g*_ and end-to-end distance *ETE*, recovered from MD simulations using all-atom (CHARMM36m) and Martini 3 models (color). The average squared ratio 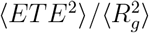 is indicated in both cases. (C) Average interresidue distance *d*_*ij*_ between the i-th and the j-th saccharide of the chain (scatterplot: av. ± s.e.). The line shows the fit ⟨*d*_*ij*_⟩ = |*b i* − *j*| ^*ν*^, where *b* is the Kuhn length (in nm) and *ν* is the scaling exponent. Resulting fitting parameters are indicated for both studied cases. (D) Jensen-Shannon divergence (JSD) of inter-saccharide distance distributions between all-atom and coarse-grained models (JSD=0: identical distributions, and JSD large: dissimilar distributions). (E) Inter-saccharide distance when |*i* − *j*| = 3 for two possible fragments, GAL-GLA-GAL and GLA-GAL-GLA (see A). C and E follow the same color scheme as B.

Polymers are also highly influenced by the surrounding solvent. To confirm that the balance between polymer-polymer and polymer-solvent interactions was maintained in our Martini 3 CSA system, we quantified the Flory scaling exponent, *ν*, from the distance *d*_*ij*_ between saccharides *i* and *j* along the CSA chain (Figure 3C). Our coarse-grained model reproduced almost perfectly the distances observed in the all-atom case. More specifically, the CSA scaling exponent obtained from distances was very close in the two descriptions, i.e. 0.65 for the all-atom and 0.66 for the coarse-grained model. From polymer theory, under equal contributions of polymer self-interactions and polymer-solvent interactions *ν* = 0.5, while *ν >* 0.5 indicates a dominance of the protein-solvent interactions rendering a highly solvated and ‘‘swollen” chain^81^. Our results indicate the latter to be the case for the studied (21 monomer long) CSA chain.

Lastly, we inspected the pairwise distances between CSA chain units by also comparing the distributions using Jensen-Shannon divergence (Figure 3D). For most distributions, JSD did not exceed 0.3 indicating a high concordance between CG and AA models. An exception to this was the pairwise distance between neighboring GAL residues, where a JSD score of 0.59 was observed. This resulted in the distance being overestimated by approximately 0.1 nm in coarse-grained model compared to the all-atom one (Figure 3E). Conversely, the JSD score for neighboring GLA residues was 0.28 indicating a good overlap between coarse-grained and all-atom distributions (Figure 3D,E).

In summary, our CG Martini model reproduces well the conformational properties and the balance between glycan-glycan and glycan-solvent interactions of a short CSA chain. Note that a comparable behavior was also observed when CG simulations were performed with original Martini 3 parameters (Figure S5B–D), but, as in the case of the disaccharide, we showed here the results for the optimized parameters that did not lead to chain aggregation, as explained in the following section.

### CSA inter-chain interactions

It is not only single chains in isolation, but also multiple coexisting chains, the topic of interest for CSA research. Accordingly, a key requirement for our coarse-grained model was to ensure that chains remain well solvated and dispersed. In order to check that, we carried out simulations of a mixture containing four 21mer chains (corresponding to a molar concentration of 1.27 mM) and monitored their aggregation tendency. The level of aggregation was quantified as the ratio between the SASA of all chains together, *S*_*tot*_, and the sum of the SASA of each chain, 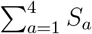. This ratio equals one if the chains are not in contact (no aggregation) and tends to zero if the chains get in contact and thus aggregate. Due to the strong electrostatic repulsion between the negatively-charged sulfate and carboxylic acid groups, at an all-atom resolution, in the microsecond time scale, the chains repelled from each other not showing any sign of aggregation (see data for CHARMM36m in Figure 4A,B). On the contrary, coarse-grained simulations using default Martini 3 parameters revealed a strong aggregation tendency (see Martini 3, Reaction-field, *q*(*ions*) = ±1.0, tiny (TQ5) case in Figure 4B). Calculation of radial distribution functions demonstrated that in this case, the sodium density around negatively charged CSA groups was found to be much higher than in the all-atom system (Figure S6A). We also noted that these cations can get sandwiched among negatively charged CSA beads, thereby bridging the distance between CSA chains (Figure S6B). Thus, we explored whether such observed inter-chain association could be prevented by reducing the strength of the electrostatic interaction between sodium and the anionic CSA moieties.

**Figure 4.**
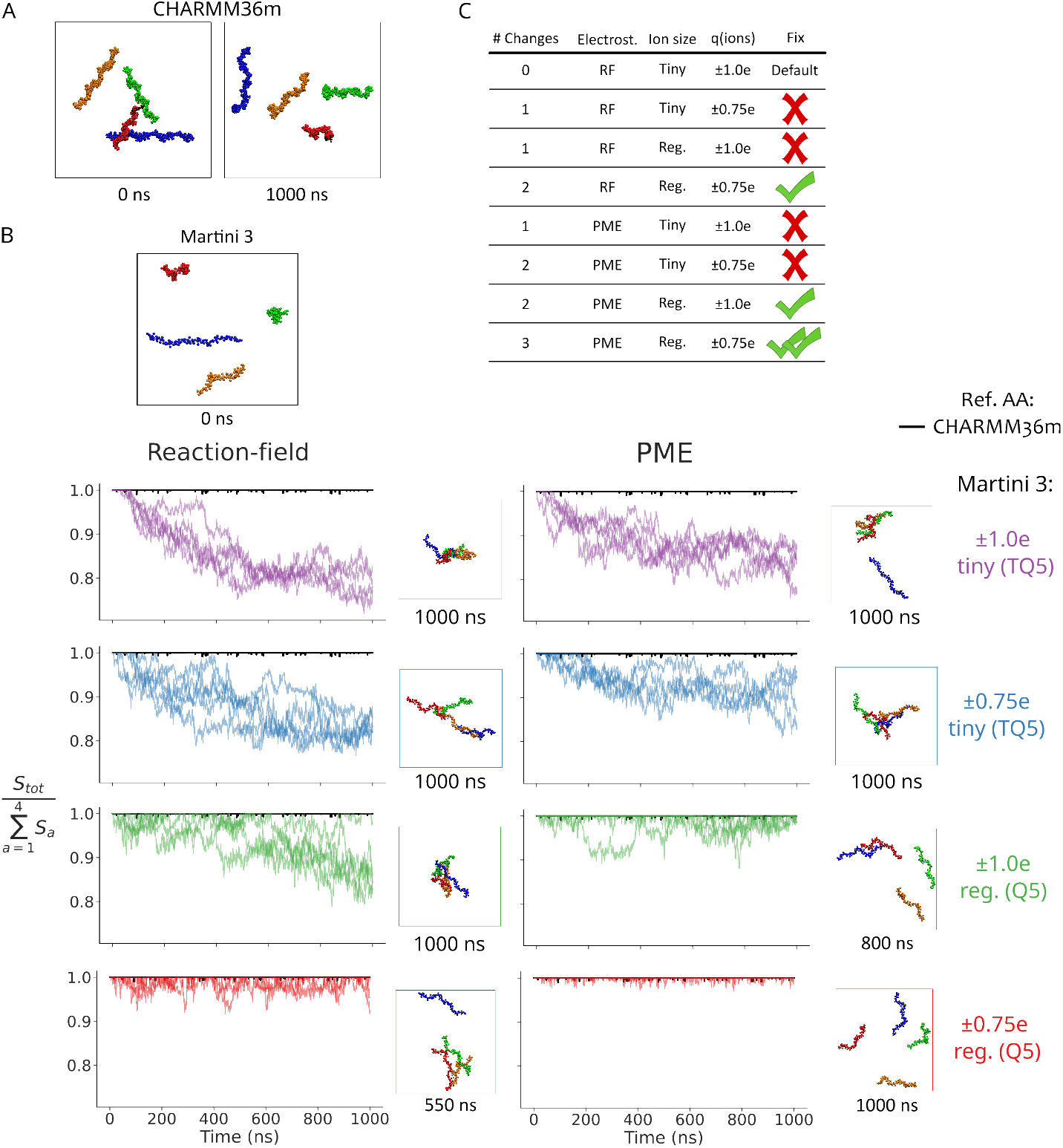
Electrostatics and ionic strength control CSA aggregation in Martini 3. (A) Snapshots extracted from all-atom (CHARMM36m) simulations of a 4xCSA 21mer system. (B) At the top, a snapshot of said system, simulated under different CG setups, is shown. At the bottom, the time traces of the chain exposure ratio, are presented (n=5). 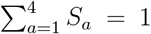 corresponds to no chain aggregation. The lower this ratio the higher the aggregation was. The electrostatic treatment (Reaction-field or PME), sodium and chloride ion bead size (tiny TQ5 or regular Q5) and the ionic charge (*q* = ± 1.0 e, or *q* = ±0.75 e) were systematically varied for Martini 3 simulations. The default condition used in Martini 3 corresponds to Reaction-field, with tiny (TQ5) sodium and chloride beads and ionic charges of *q* = ±1.0*e* (top-left panel, purple). Next to the time traces a snapshot from each simulation at the indicated time is shown. Time-traces for atomistic simulations are shown in black as reference. In this case the exposure ratio was almost 1, in the studied time-scale, indicative of no aggregation, (C) Summary of studied conditions, indicating to which extent they fixed aggregation. Individual CSA chains in simulations snapshots are shown in different colors (red, blue, green, yellow).

Three aspects of relevance for the interaction of glycans with ions were considered. Firstly, the Coulomb potential in Martini 3 is usually handled using the reaction-field scheme ^48^. Although being less computationally intensive, this method can lead to artifacts, including truncation and discontinuities of the potential function and a mean-field approximation of long range electrostatics^85^. Thus, we tested Particle Mesh Ewald (PME), as an alternative. PME alleviates these issues at the expense of a higher computational cost. PME is the standard in all-atom simulations although it has also been used in Martini coarse-grained simulations, e.g of an ionic liquid^86^. Secondly, a hydration shell forms around ions suspended in water. In older versions of Martini, this was implicitly accounted for by representing ions with bead sizes larger than their ionic radius^87^. However, this practice was abandoned in Martini 3, resorting to the use of the smallest bead TQ5. The choice of bead radius would not only influence the size of the ion particle but also affect its Lenard-Jones potential parameters. For this reason, we explored the effect of using a larger bead for monovalent ions (i.e. Q5). Lastly, it has been suggested that electrostatic interactions tend to be largely overestimated in molecular dynamics simulations with non-polarizable force-fields^88^. Authors in ref.^88^ proposed charge rescaling proportional to electronic polarization contribution of water screening as a way to more accurately capture Coulombic forces. We included such a correction by reducing the charge of monovalent salt ions from ±1.0 to ±0.75 (both for sodium and chloride). Consequently, we conducted equilibrium simulations of the 21-mer CSA chain mixture with all possible combinations of these three conditions, to test how they influenced inter-chain association.

As mentioned, the default Martini 3 simulations setup (i.e. RF treatment, tiny TQ5 beads of *q* ± 1.0 e for sodium and chloride) had the largest decrease in the exposed area ratio, implying the largest level of aggregation (Figure 4B, top-left). Changing each of the three above-mentioned aspects individually reduced aggregation, however not fully (Figure 4B): for RF versus PME compare left-column and right-column purple curves, for bead size compare the tiny bead (purple) and the regular bead (green) curves, and for the charge scaling compare *q* = ±1.0*e* (purple) with *q* = ±0.75*e* (blue). The situation improved when two conditions were adjusted at the same time. In two simulation setups (either RF with regular-bead size (Q5) and rescaled-charge or PME with regular-bead size (Q5) and full *q* = ±1.0*e* charge) the exposure ratio fluctuated around one, indicating the formation of brief and unstable small aggregates (Figure 4B). Interestingly, using PME and rescaling the ionic charges, while still describing sodium with a tiny bead, did not show a significant improvement. Finally, aggregation was fully avoided, as observed in all-atom simulations, when all three conditions, i.e. PME, (Q5) regular-size beads, and rescaled ionic charge of *q* = ±0.75*e*, were in place (Figure 4B).

In summary, we identify at least three distinct possible treatments of glycan-ion electrostatic interactions that give satisfactory aggregation levels of CSA at a coarse-grained level of resolution (Figure 4C). To illustrate the applicability of our coarse-grained model in two relevant contexts, in the following we consider the conditions that led to full abolishment of aggregation, i.e. after using PME and regular-size beads for sodium or chloride ions with rescaled charges.

### CSA in complex with VAR2CSA

After establishing a coarse-grained model for CSA and non-aggregating simulation conditions, we sought to confirm that such setup can accurately capture dynamics of a protein in complex with the glycan. To test this, we have chosen the malaria adhesin VAR2CSA which is known to bind CSA chain in its fully sulphated form^23,24,89^. The interaction of VAR2CSA with CSA is essential for attachment of malaria-infected erythrocytes to the placenta proteoglycan matrix, leading to pregnancy associated malaria^90^. Hence, disruption of this interactions is a promising therapeutic approach for the design of anti-malarial vaccines^91^. This complex has previously been investigated with all-atom simulations^38,40^. Thus, we here attempted to reproduce the (all-atom) equilibrium dynamics of this complex using our Martini 3 model for CSA. The coarse-grained model of VAR2CSA was generated following standard Martini 3 procedure by applying a Gō-type additional interactions (see methods)^62^. The initial conformation of the complex at both levels of resolution is illustrated in Figure 5A. The dynamics of the complex were then simulated in *n* = 10 independent replicas of 300 ns each.

**Figure 5.**
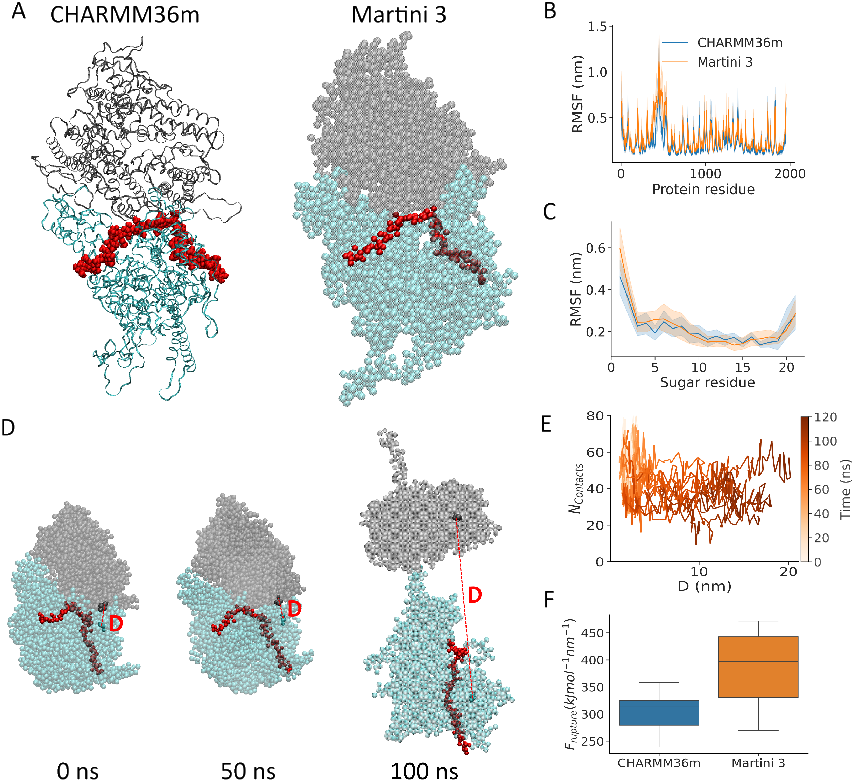
Coarse-grained simulations of CSA bound to the malaria adhesin VAR2CSA. (A) Snapshot of VAR2CSA bound to CSA 21-mer (red) in all-atom and corse-grained Martini 3 resolution. VAR2CSA N-terminal subunit (residues 1-964) is marked in cyan and C-terminal subunit (residues 965-1954) -in gray. (B,C) Root mean square fluctuation (RMSF) of protein backbone (B) and the CSA sugar chain (C) obtained from equilibrium MD simulations. All-atom simulations taken from.^40^ (D-F) Coarse-grained pulling simulations of VAR2CSA. Representative snapshots during the opening process are depicted in D (same color code as in A). Residues used to measure distance between N and C terminal subunits (red dashed line) were 120 and 1663 respectively (opaque). (E) Number of contacts between VAR2CSA and CSA as distance between the N- and the C-terminal subunits changes. (F) Force needed for unfolding. B,C,E,F show data from n=10 independent replicates.

As expected, by imposing Gō distance-restraints, the protein largely retained its structural fold (Figure S7). Local dynamics were subsequently compared using per residue backbone root-mean-square fluctuations (RMSFs) (Figure 5B). There was a good agreement of residue flexibility, with only deviations arising from highly mobile regions. When monitoring the local flexibility of the (bound) CSA chain, also via RMSF, we also observed good agreement between all-atom and coarse-grained simulations (Figure 5C). Saccharides 3 to 19 were directly in contact with VAR2CSA and thus displayed reduced flexibility, compared to chain ends which were not in contact and hence more mobile. The high agreement of RMSF values demonstrate that the equilibrium dynamics of CSA in complex with a protein can be accurately reproduced by our glycan model in conjunction with a conventional protein model used for Martini.

Previous atomistic simulations have also demonstrated that applying elongational tension to VAR2CSA causes an opening of the core region^40^, a process of relevance for the shear-enhanced adherence mechanism of malaria-infected erythrocytes mediated by this protein^92^. We thus checked if our CG model was also capable to capture the force response of this complex. By performing constant-velocity force-probe MD simulations, applying the same elongational tension on the C-terminal methionine and 21’st sugar residue of the CSA chain we were able to achieve the same type of opening behavior (Figure 5D). The distance between the two subdomain regions suddenly increased upon surpassing a certain force threshold (*F*_*rupture*_) value and the opening always preceded glycan dissociation (Figure 5E). We quantified the rupture force by identifying the timepoint when the VAR2CSA inter-domain distance was ≥ 5 nm and then identifying the maximal pulling force within 10 ns window around this timepoint. *F*_*rupture*_ in CG simulations was greater than in all-atom simulations, likely because default Gō restraints were acting on the opened interface creating additional resistance not present in atomistic protein model (Figure 5F). However, the difference was not substantial and in agreement with all-atom simulations, the force required to open the VAR2CSA core region was lower than the force needed for CSA dissociation and thus opening always preceded dissociation(Figure 5E, F). Thus, not only under equilibrium, but also under non-equilibrium tension conditions, our coarse-grained model of CSA (together with a Gō-type protein description) properly captures the specific properties of the VAR2CSA-CSA complex.

### Simulations of a CSA 123mer

So far we have focused our analysis on 21mer CSA chains to allow comparison with allatom simulations and demonstrated that our coarse-grained model accurately reproduces the dynamics of atomistic CHARMM36m simulations. However, *in vivo* CSA chains are often longer than 100 saccharides - a scale which is no longer feasible to simulate at an all-atom resolution. Therefore, we proceeded to simulate a CSA 123-mer by using our CG Martini 3 model for 10 independent replicates of 2 *µs* each (Figure 6A). Surprisingly, although the energy surface of the coarse-grained resolution was smoother, these simulations revealed slow equilibrium dynamics, as reflected by slow-taking gradual fluctuations of the end-to-end distance (Figure S8). Thus, not only does the Martini 3 model of CSA enabled simulations of more biologically relevant size scales, it also allowed us to capture conformational states that would take much longer to sample atomistically.

**Figure 6.**
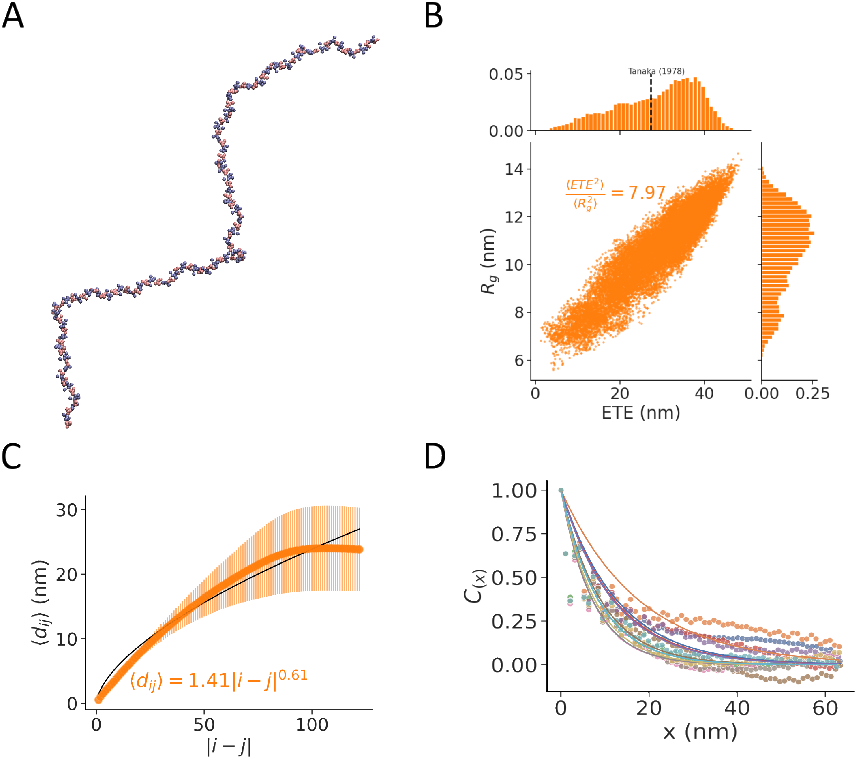
Coarse-grained CSA 123mer simulations. (A) Snapshot of 123-sugar CSA chain at CG resolution (GLA: red and GAL: blue). (B) Ratio between radius of gyration *R*_*g*_ and end-to-end distance *ETE* and their distributions, recovered from MD simulations. Text indicates the average squared ratio 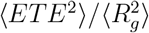. Dashed line indicates experimental mean root squared end-to-end distance^13^. (C) Average inter-residue distance ⟨*d*_*ij*_⟩ between the i-th and the j-th monomer of the chain (scatterplot: av. ±s.e.). Black line shows the fit ⟨*d*_*ij*_⟩ = *b*| *i* − *j*| ^*ν*^, where *b* is the Kuhn length, 1.41 nm, and *ν* is the scaling exponent, 0.61. Decay of autocorrelation C(x) of GLA virtual sites with distance x (simulation data: points and exponential decay fit: lines). Distance at which C(x)=0.5 was considered the persistence length. Different colors represent *n* = 10 independent replicates.

The simulations of the 123mer allowed us for quantify polymer properties of a fully sulfated CSA chain in explicit solvent. Consistent with the CSA 21mer, we find that the ratio 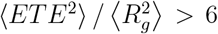 pointing to an extended conformation (Figure 6B). In addition, the mean end-to-end distance of this chain was 28.7 nm which is close to the experimentally measured value for a chain of similar molecular weight of 27.3 nm^13^, while previous estimates with an implicit-solvent CG model^43^ predicted values of the order of 32.4 nm. The scaling exponent is similar to our estimate for a CSA 21-mer, confirming the dominance of polymer-solvent interactions (Figure 6C). In addition, the characteristic ratio, i.e. 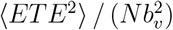 (with *N* = 123, the polymerization degree, and *b*_*v*_ = 0.52 nm, the bond length), was found do be 27.6, which falls in the same range of a previous estimate from an implicit-solvent CG model (between ∼30–35 for a comparable *N* in Figure 6 in Bathe et al.^43^). Previous studies using implicit-solvent CG simulations and dynamic light scattering experiments reported hydrodynamic radii, *R*_*h*_, values of around 5.5 nm, for CS chains of comparable polymerization degree^45^ while explicit-solvent CG simulations (using Martini 2.2) reported values of ∼ 4.8 ± 0.3 nm for a 100-monomer fully-solvated chain^47^. Assuming a linear monodisperse chain, *R*_*g*_ = 1.5*R*_*h*_ ^45,93^, our estimate of the radius of gyration, *R*_*g*_ = 10.6±1.7 nm (mean±s.d., Figure 6B), turns into a *R*_*h*_ = 7.1 ± 1.1 nm. Although larger, our estimate is not too far from such previous estimates, specially taking into account the differences in the samples (i.e. size and sulfation distribution), environmental conditions and the approximate nature of the *R*_*g*_ to *R*_*h*_ conversion.

Lastly, we estimated the persistence length to vary between 6.3 nm and 17.8 nm, with a mean of 9.8 nm (Figure 6D), the large variation attributed to the slow dynamics of the chain. Encouragingly, our values are not far from previous computational persistence length estimates, which saturated from 11.2–12.0 nm for several hundreds, i.e. N=256, chains^43^, and from a small angle neutron scattering estimate of 12.7 nm (for a 0.01 w/w concentration)^94^.

## Discussion

Chondroitin sulfate A (CSA) is a linear glycosaminoglycan that is attached to cell surface or secreted proteins^1^. It regulates humoral signaling pathways^17^, ensures hydration and toughness of various tissues^10–12^, acts as an antioxidant^15^ and is implicated in many other vital roles. Research into CSA has been difficult due to its large size, heterogeneous chemical composition and multitude of binding partners. Here, we developed a model of CSA for coarse-grained simulations based on the Martini 3 force-field to address these challenges.

Previous high-resolution simulations of CSA have been limited to short fragments^31–33,35^. In our CG model, the 53 atoms constituting the disaccharide repeating unit are represented by 11 Martini particles. Despite of this reduction in the level of detail, our model still retained an accurate representation of the repeating-unit’s molecular volume and its internal bonded interactions (Figure 2). Subsequently, the parameters of the repeating unit could be used to build longer CSA chains. For medium-sized chains, of a couple tens of monomers, our results compare extremely well to atomistic simulations (performed using the CHARMM36m force-field^51^) in terms of description of polymeric properties, such as the chain extension and inter-monomer interactions (Figure 3). The minimal CSA segment that binds to the malarial adhesin VAR2CSA has such a degree of polymerization^24^ and we thus show that our CG model is capable of capturing the key structural details of CSA fragments of that length.

Beyond short fragments, the modularity of the model allowed to investigate the behavior of much longer CSA chains. The degree of polymerization of this glycan in nature is of the order of several tens or even hundreds of monomers^95^. For a chain in that range (exactly 123-monomers), we predict structural properties that are encouragingly close to experimental estimates^13,45,94^ and to previous estimates from implicit-solvent^43,45^ and explicit-solvent^47^ CG models (Figure 6). Although it should be noted that differences in the exact size and sulfation distribution, as well as in buffer conditions, make this comparison not straightforward. In particular we predict such chain to adopt extended conformations beyond what it would be expected for a Gaussian chain^84^ (Figure 6B), also reflected by the markedly long persistence length of the order of 10 nm (20 monomers) (Figure 6D). This is likely due to strong self-repelling electrostatic forces, promoting solvent-glycan interactions over glycan-glycan ones (Figure 6C). Note that such estimate could not have been obtained from the simulations of short fragments of size comparable to the persistence length.

When solvated by explicit Martini water beads and ions, the 123mer CSA chain system consisted of almost 1 million CG beads. One microsecond of molecular dynamics of such a system on a 2 GPU and 72 CPU node required 7 days of computing. If treated atomistically, such simulation would have implied roughly 11 million atoms. Although Martini CG and all-atom time-scales are not directly comparable^87^, microsecond-long simulations of such system would be extremely time and resource-consuming, i.e. require approximately 500 days of computing using a single 128 CPU node^96^. We thus provide evidence how our model enables the access to spatial and temporal scales that are currently extremely difficult with atomistic simulations.

Despite accurately capturing single-chain properties, simulations of multiple chains, using the recommended Martini 3 interaction and simulation parameters resulted in aggregation (Figure 4). We attribute this tendency to the strong electrostatic interaction between the negatively charged sugar groups and sodium ions, that promoted inter-chain association of the CSA chains bridged by sodium ions (Figure S6). We systematically varied three key electrostatic properties of the system that attenuated the strong interactions with sodium and quantified their impact on the aggregation of CSA (Figure 4B). By using reaction field scheme, we demonstrated that aggregation could be prevented by both increasing the size of ion beads, reflecting their hydration shell (as it was originally implemented in previous Martini version^87^), and rescaling charge of salt ions, according to previously proposed continuum correction to take into account electrostatic screening of water^88^. Charge scaling is actually preferable to directly changing the screening constant, as it does not affect all other Coulombic interactions of the system. Note that both corrections had to be in place in order to prevent aggregation. By using PME, which is a more robust way to consider long-range electrostatic interactions, the extent of aggregation was overall reduced but not fully abolished, requiring a bigger (hydrated) ion bead too. Furthermore, if the three conditions were in place, i.e. PME and bigger salt ion beads with rescaled charges the CG model did not show aggregation, similarly as the atomistic reference. Note that aggregation has been reported in previous simulations with Martini (version 2.2) ^47,97,98^. Rather than rescaling Lennard-Jones parameters, as previously proposed for polysaccharides^47,98^, we here propose tuning electrostatic interactions as an approach to mitigate it. Accordingly, we provide at least three schemes of varying level of modifications (in reference to default Martini 3 force field) to attenuate glycan-cation interactions and thereby prevent CSA aggregation (Figure 4C).

Treatment of electrostatics of highly charged systems is a challenging task both in allatom^88^ and coarse-grained MD simulations. Because CSA is a highly sulfated and thus highly negatively-charged biopolymer, studying its aggregation becomes an excellent test to assess electrostatics in MD simulations. Already at the all-atom level, electrostatics that govern CSA interaction with ions have been found difficult to reproduce and to be highly sensitive to force-field parameters^33^. At the coarse-grained level,stiffness of CSA chains^43^. At that level of resolution, we here demonstrate the impact of long-range electrostatics (reaction-field vs PME), ion solvation (small vs. big CG ion beads), and electric screening (fractional ionic charges) have on glycan clustering. This information provides some practical clues for future studies, aiming at solving the issues existing with cations in Martini, not only for the simulation of glycans, but more generally for studying highly-charged systems.

We based our model in the Martini 3 force-field as it enables simulations of large and complex multi-component systems in coarse-grained resolution. This allowed us to simulate multiple (Figure 4) and also large (Figure 6) CSA chains in explicit solvent. We thereby complement existing Martini models of Glycosaminoglycans ^47^, available for the 2.2 version, by providing here a model of CSA, the most common CS form, compatible with the Martini 3 version. Considering the solvent explicitly may constitute an advantage over existing implicit solvent models of CSA^43–46^, when evaluating solvent properties such as viscosity or friction. This will be of relevance to study processes such as lubrication, load bearing of joints, or shear-mediated cellular adherence. In addition, our model can be used in conjunction with models of other biomolecules such as proteins. We demonstrate this by simulating VAR2CSA-CSA complex (Figure 5). This constitutes another advantage of our model, as biologically-relevant questions related to interactions of CSA with other biomolecules can now be addressed within the Martini 3 framework.

In conclusion, we here have developed a Martini 3 coarse-grained model of chondroitin sulfate A that allows its MD simulations at reduced computational cost without compromising simulation accuracy. This opens the door for exciting research to study CSA and potentially other chondroitin sulfates.

## Acknowledgement

This work was supported by the Klaus Tschira Foundation (to all authors). This research was conducted within the Max Planck School Matter to Life supported by the German Federal Ministry of Education and Research (BMBF) in collaboration with the Max Planck Society. Computations were performed on the HPC system at the Max Planck Computing and Data Facility.

## Supplementary Figures

**Supplementary Figure 1.**
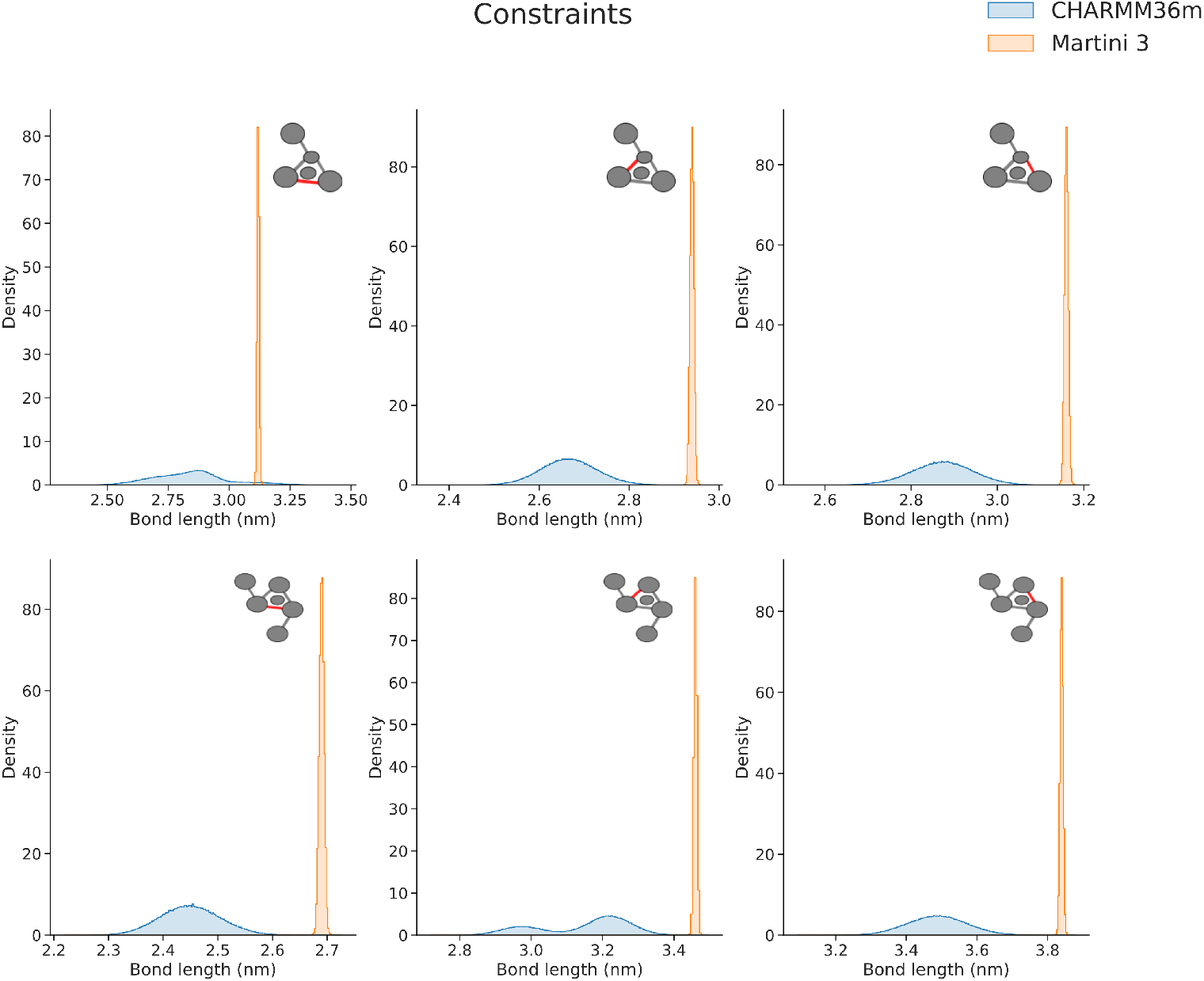
Ring bonds are represented as distance constraints in Martini 3 model. Distance comparison between forward mapped all-atom trajectory (CHARMM36m) and Martini 3. Inlet shows the corresponding bond marked in red.

**Supplementary Figure 2.**
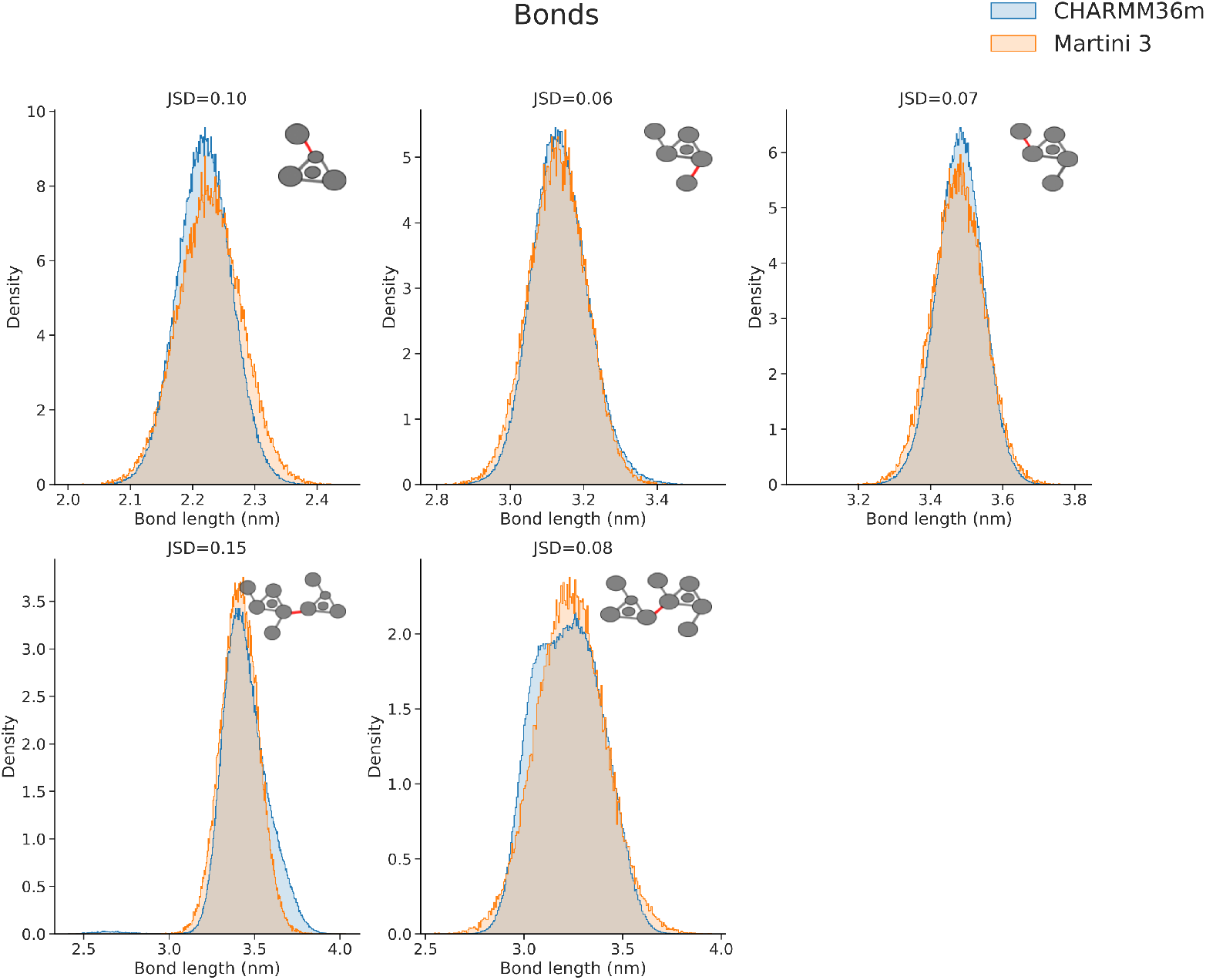
Comparison of bond length distributions. Bond length comparison between forward mapped all-atom trajectory (CHARMM36m) and Martini 3. Inlet shows the corresponding bond marked in red.

**Supplementary Figure 3.**
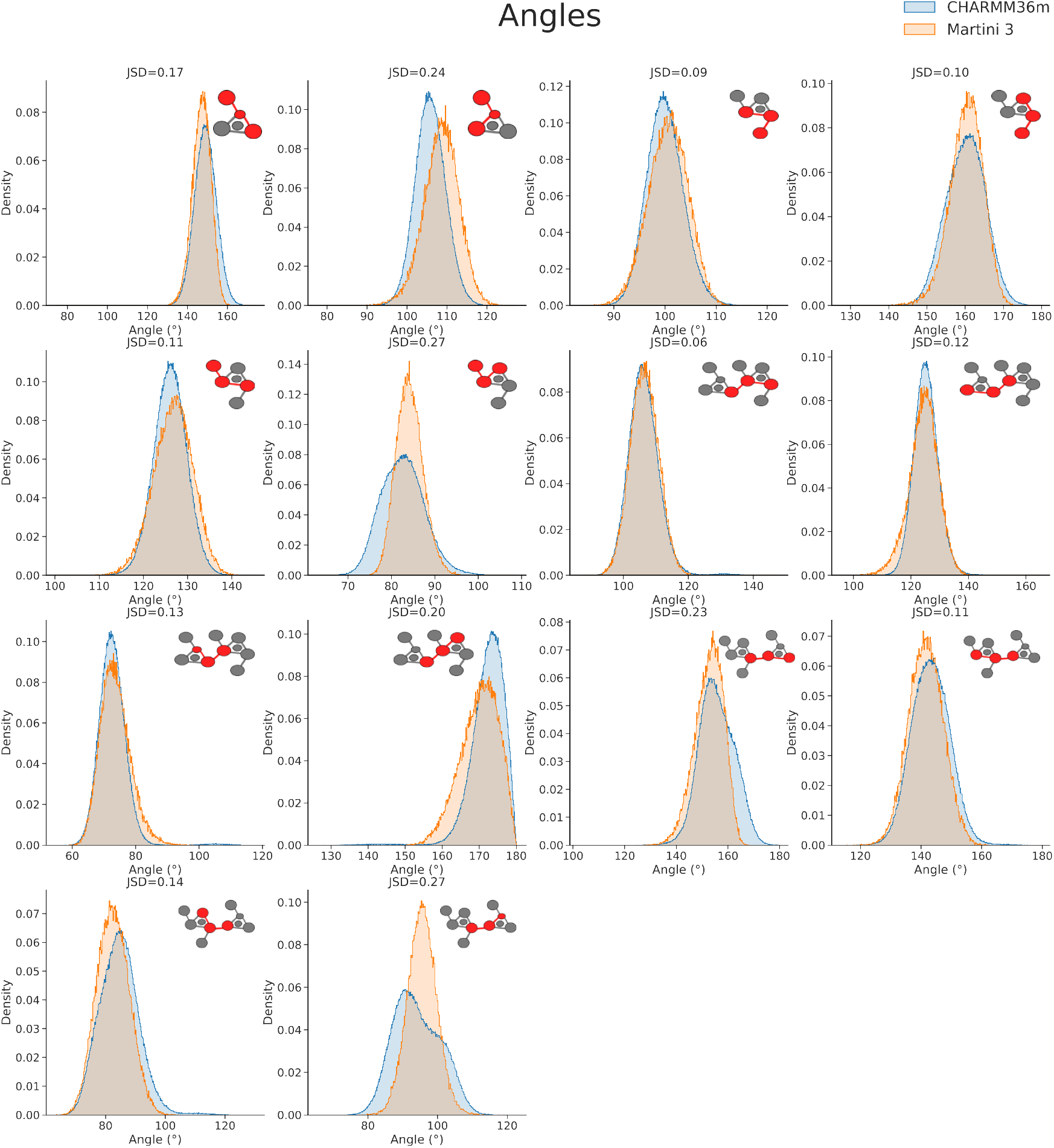
Comparison of angle distributions. Angle comparison between forward mapped all-atom trajectory (CHARMM36m) and Martini 3. Inlet shows the corresponding angle marked in red.

**Supplementary Figure 4.**
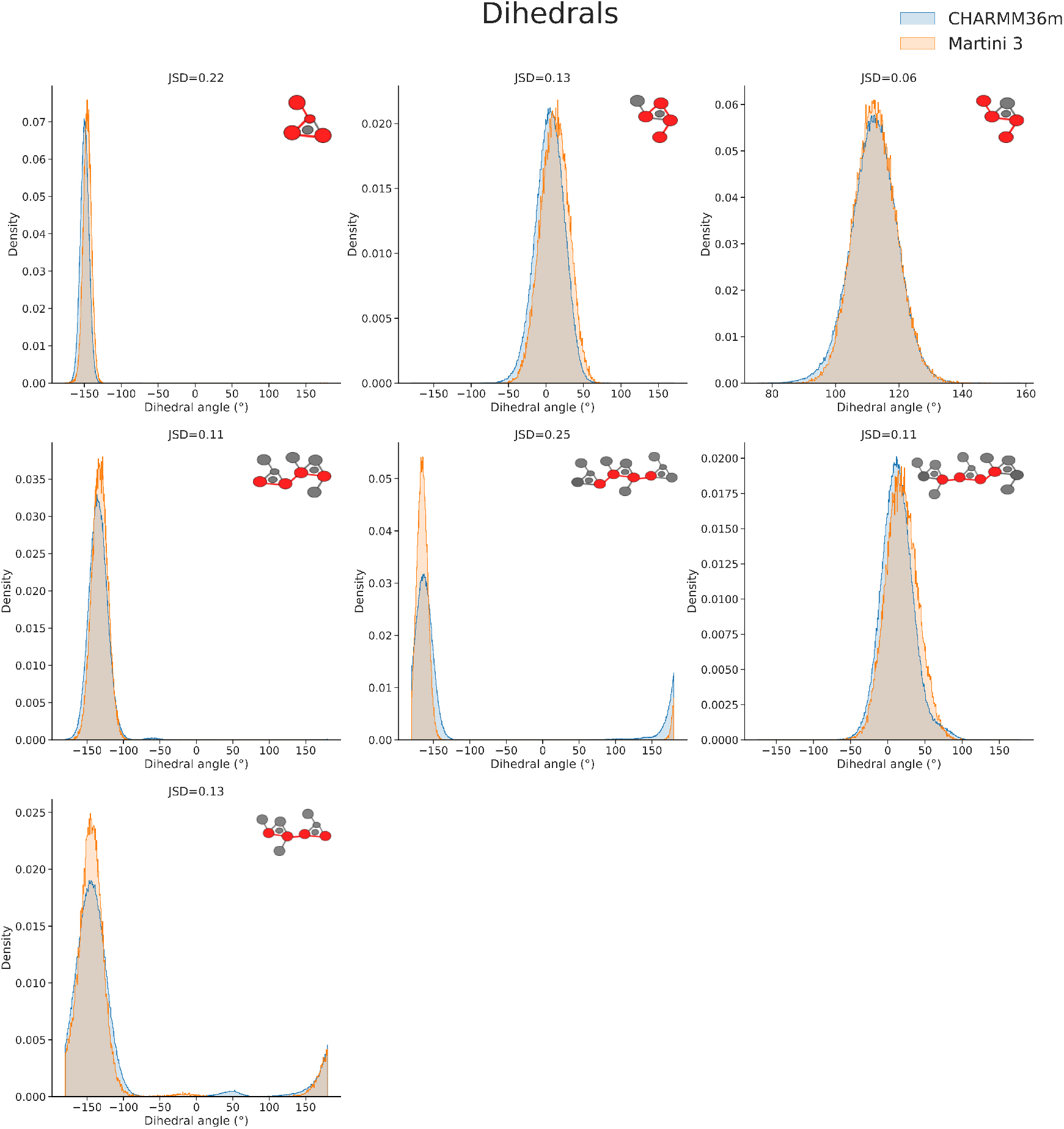
Comparison of dihedral angle distributions. Dihedral angle comparison between forward mapped all-atom trajectory (CHARMM36m) and Martini 3. Inlet shows the corresponding dihedral angle marked in red.

**Supplementary Figure 5.**
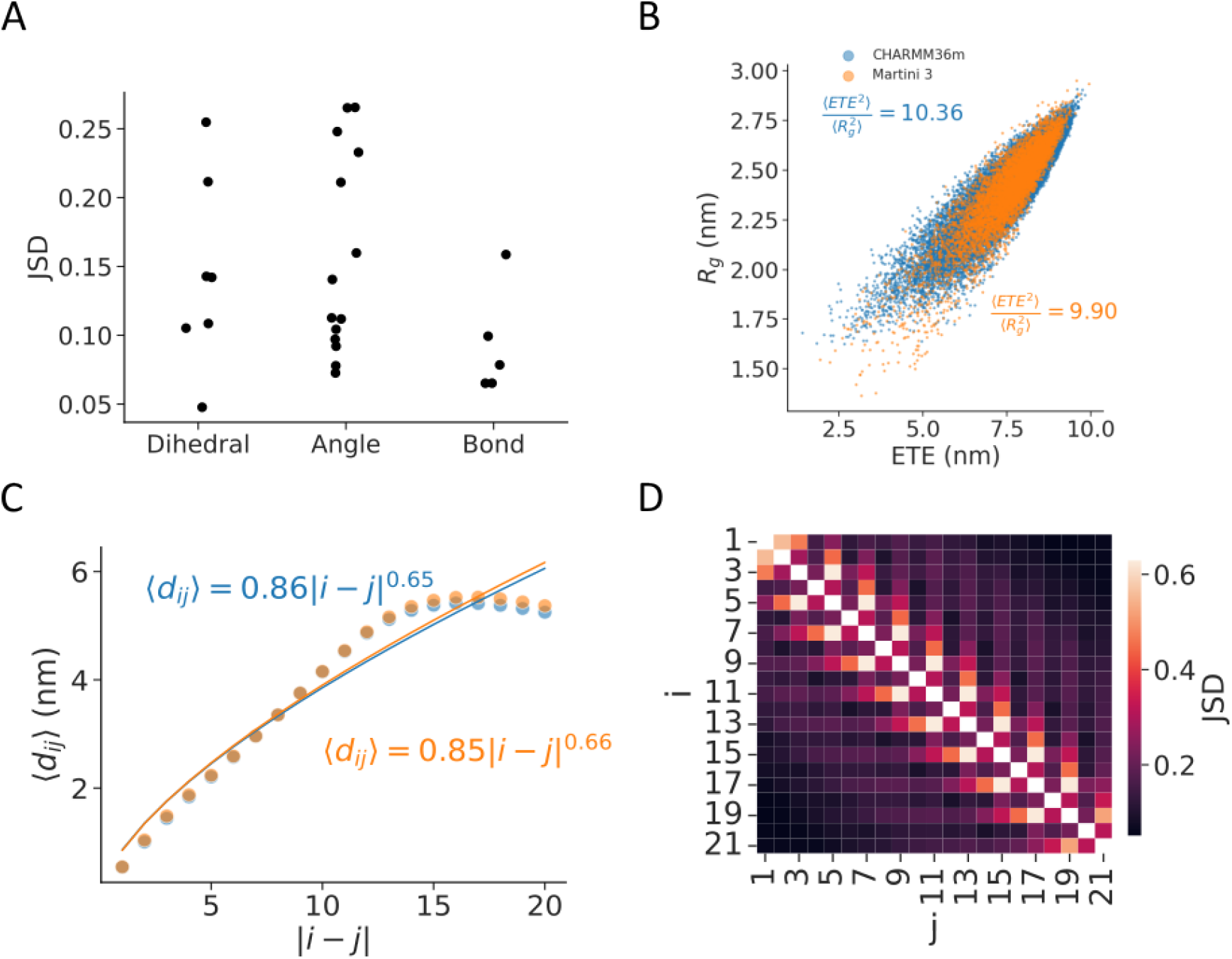
Performance of CG CSA simulation with default Martini3 parameters. (A) Jensen-Shannon divergence statistic for the three types of bonded terms used, comparing the Martini 3 model with the all-atom CHARMM36m reference data (JSD=0: identical distributions, and JSD large: dissimilar distributions). (B) Ratio between radius of gyration *R*_*g*_ and end-to-end distance *ETE*, recovered from MD simulations using all-atom (CHARMM36m) and Martini 3 models (color). The average squared ratio 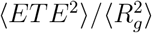 is indicated in both cases. (C) Average interresidue distance ⟨*d*_*ij*_⟩ between the i-th and the j-th monomer of the chain (scatterplot: av. ±s.e.). The line shows the fit ⟨*d*_*ij*_⟩ = *b* |*i* − *j*| ^*ν*^, where *b* is Kuhn length in nm and *ν* is the scaling exponent. Resulting fitting parameters are indicated for both studied cases. Colors follow the same scheme as in B (D) JS divergence of interresidue distance distributions between all-atom and coarse-grained models.

**Supplementary Figure 6.**
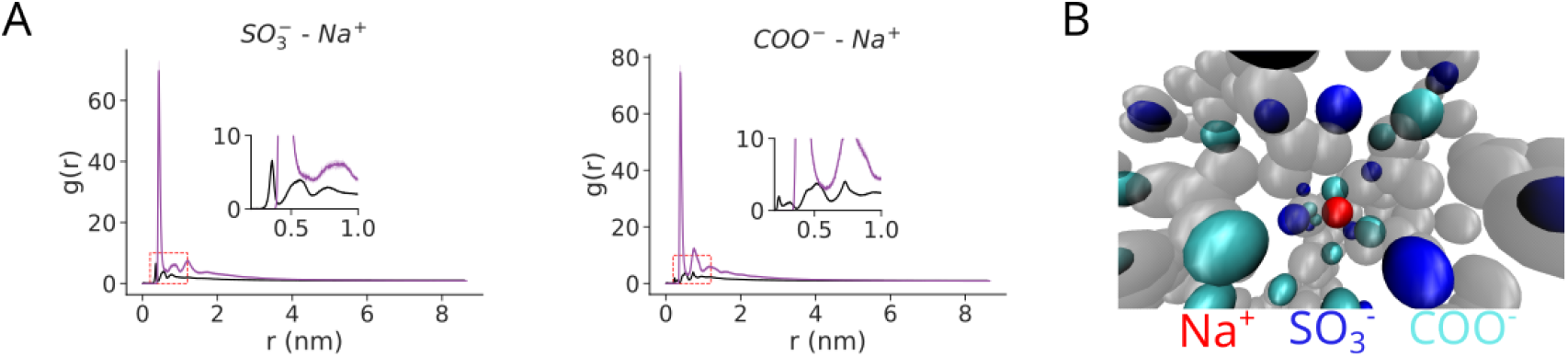
Sodium cation interactions are overestimated in coarsegrained simulations. (A) Comparison of radial distribution functions of sodium ions around carboxylic acid and sulfate groups. Inlet shows a zoom on the region marked by the red rectangle. Lines follow the same coloring as Figure 4A. (B) Snapshot of sodium cation (red) sandwiched inside CSA aggregate (gray), between negatively charged sulfate (blue) and carboxylic acid (cyan) beads.

**Supplementary Figure 7.**
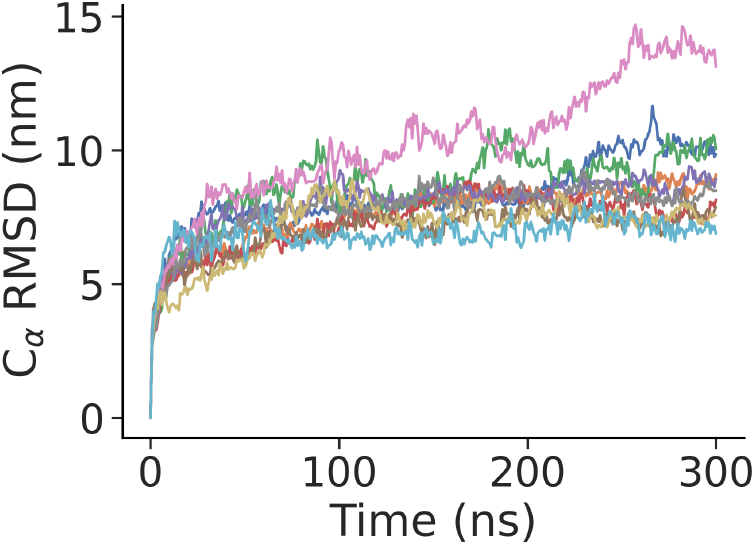
*C*_*α*_ root mean square deviation (RMSD) of VAR2CSA coarse-grained model. Different colors show n=10 independent replicates.

**Supplementary Figure 8.**
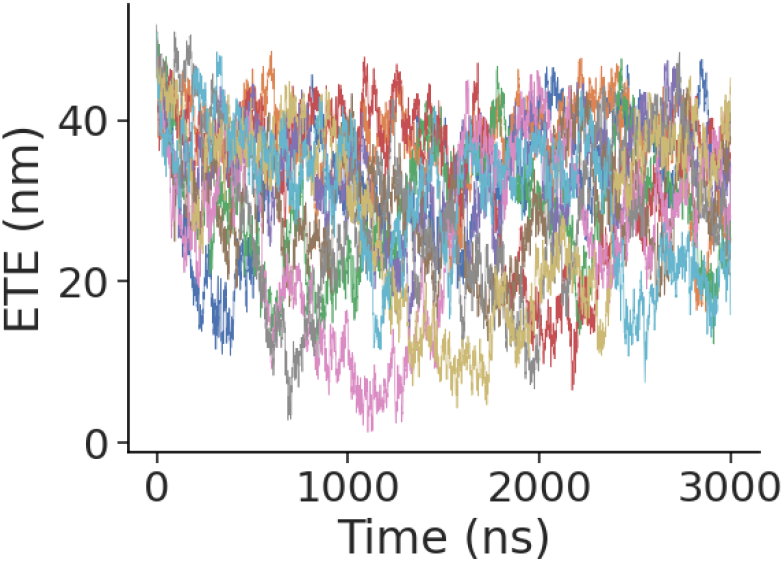
Time traces of end-to-end distance fluctuations of CSA 123mer. Different colors represent individual simulation replicates.

